# Combination Respiratory Vaccine Containing Recombinant SARS-CoV-2 Spike and Quadrivalent Seasonal Influenza Hemagglutinin Nanoparticles with Matrix-M Adjuvant

**DOI:** 10.1101/2021.05.05.442782

**Authors:** Michael J Massare, Nita Patel, Bin Zhou, Sonia Maciejewski, Rhonda Flores, Mimi Guebre-Xabier, Jing-Hui Tian, Alyse D. Portnoff, Louis Fries, Vivek Shinde, Larry Ellingsworth, Gregory Glenn, Gale Smith

## Abstract

The 2019 outbreak of a severe respiratory disease caused by an emerging coronavirus, SARS-CoV-2, has spread globally with high morbidity and mortality. Co-circulating seasonal influenza has greatly diminished recently, but expected to return with novel strains emerging, thus requiring annual strain adjustments. We have developed a recombinant hemagglutinin (HA) quadrivalent nanoparticle influenza vaccine (qNIV) produced using an established recombinant insect cell expression system to produce nanoparticles. Influenza qNIV adjuvanted with Matrix-M was well-tolerated and induced robust antibody and cellular responses, notably against both homologous and drifted A/H3N2 viruses in Phase 1, 2, and 3 trials. We also developed a full-length SARS-CoV-2 spike protein vaccine which is stable in the prefusion conformation (NVX-CoV2373) using the same platform technology. In phase 3 clinical trials, NVX-CoV2373 is highly immunogenic and protective against the prototype strain and B.1.1.7 variant. Here we describe the immunogenicity and efficacy of a combination quadrivalent seasonal flu and COVID-19 vaccine (qNIV/CoV2373) in ferret and hamster models. The combination qNIV/CoV2373 vaccine produces high titer influenza hemagglutination inhibiting (HAI) and neutralizing antibodies against influenza A and B strains. The combination vaccine also elicited antibodies that block SARS-CoV-2 spike protein binding to the human angiotensin converting enzyme-2 (hACE2) receptor. Significantly, hamsters immunized with qNIV/CoV2373 vaccine and challenged with SARS-CoV-2 were protected against weight loss and were free of replicating SARS-CoV-2 in the upper and lower respiratory tract with no evidence of viral pneumonia. This study supports evaluation of qNIV/CoV2373 combination vaccine as a preventive measure for seasonal influenza and CoVID-19.

**Highlights:**

- Combination qNIV/CoV2373 vaccine induced protective hemagglutination inhibition (HAI) responses to seasonal influenza A and B unchanged when formulated with recombinant spike.
- Combination qNIV/CoV2373 vaccine maintained clinical and virologic protection against experimental challenge with SARS-CoV-2.
- Combination qNIV/CoV2373 vaccine showed no clinical or histological sign of enhanced disease following experimental challenge with SARS-CoV-2.
- Combination qNIV/CoV2373 vaccine induced antibodies against SARS-CoV-2 neutralizing epitopes common between US-WA and B.1.352 variant.

## 1. Introduction

The remarkable development of SARS CoV-2 vaccines in the past 12 months appears to be stemming the epidemic in geographies where implementation has been successful [1]. Concurrently, there has been notable reduction in seasonal influenza cases globally during the SARS CoV-2 pandemic, but this is widely expected to revert to rates previously experienced, with the concomitant antigenic drift in viral emerging strains [2, 3]. Co-circulating seasonal influenza virus thus presents a continued threat with the appearance of new seasonal strains and the potential for novel influenza strains periodically emerge creating a new pandemic. Seasonal influenza epidemics are responsible for 389,000 (range 294,000 to 518,000) influenza-associated respiratory deaths annually [4]. Over the past 100 years, four major influenza pandemics with high mortality have occurred -1918-1919 H1N1 Spanish flu with 50 million deaths [5], 1957-1959 H2N2 Asian flu with 1.5 million deaths [6], 1968-1969 H3N2 Hong Kong flu with 1 million deaths [7], and 2009-2010 H1N1pdm09 swine flu with 151,700-575,400 deaths worldwide [8].

We describe here the development of a novel combination respiratory vaccine consisting of a recombinant hemagglutinin (HA) quadrivalent nanoparticle influenza vaccine (qNIV) formulated together with a recombinant SARS-CoV-2 spike protein vaccine and adjuvanted with saponin Matrix-M™. Matrix-M adjuvanted quadrivalent seasonal influenza HA vaccine and adjuvanted SARS-Cov-2 spike protein vaccines recently, independently, and successfully evaluated in pivotal Phase 3 clinical trials [9-11].

We have previously shown influenza hemagglutinin (HA) and respiratory syncytial virus fusion (F) full-length nanoparticle vaccines with Matrix-M adjuvant presents conserved epitopes that induce broadly neutralizing antibodies [12-14]. In multiple clinical trials, quadrivalent seasonal HA nanoparticle influenza vaccine (qNIV) stimulates broadly functional immunity against historic and forward drift H3N2 strains. Highly conserved neutralizing epitopes in the HA stem and vestigial esterase domain were identified and may be responsible for broad functional immunity [12, 13]. Influenza qNIV induces hemagglutination inhibition (HAI) and microneutralizing (MN) antibodies against homologous and drifted A/H3N2, A/H1N1, and B strains in preclinical models [12, 13] and human clinical studies [15, 16].

Using the recombinant insect cell production platform to produce nanoparticles, a COVID-19 vaccine, NVX-CoV2373, developed based upon the full-length SARS-CoV-2 spike (S) glycoprotein that stabilized in the prefusion conformation by mutation of the furin cleavage site and introduction of two proline substitutions in the apex of the central helix [17-18]. NVX-CoV2373 nanoparticles are composed of prefusion S-trimers that associate with polysorbate 80 (PS 80) micelles to form one to multi-S trimer rosettes [19]. NVX-CoV2373 elicits high titer anti-S antibody that blocks spike binding to the human angiotensin converting enzyme-2 (hACE2) receptor and antibodies that neutralize prototype SARS-CoV-2 and newly evolved variants. Mice, cynomolgus macaques, and rhesus macaques immunized with NVX-CoV2373 vaccine were protected against upper and lower respiratory tract infection following SARS-CoV-2 challenge [17, 20, 21]. In multiple clinical trials (NCT04368988, NCT04533399, NCT04583995, NCT04611802), NVX-CoV2373 vaccine was well tolerated and immunogenic, inducing virus neutralizing antibodies and Th1-biased CD4+ T cell responses [22]; high efficacy against prototype Wuhan-Hu-1-like virus USA-WA1 and B.1.1.7 and B.1.351 variants was shown [10, 11]. Here we investigate the immunogenicity and efficacy of a combination quadrivalent seasonal influenza (qNIV) and COVID-19 NVX-CoV2373 with Matrix-M vaccine (qNIV/CoV2373) in ferrets and hamsters. We show a combination vaccine elicits undiminished protective levels of influenza A and B strain hemagglutination inhibition (HAI) and neutralizing antibodies, and protection against upper and lower respiratory infection with no evidence of viral pneumonia following challenge with SARS-CoV-2.

## 2. Materials and methods

### 2.1. Virus stock and receptor

The SARS-CoV-2 (strain 2019-nCoV/USA-WA1/2020) isolate was obtained from the Center for Disease Control and stock virus prepared by passage in Vero E6 cells. A/Kansas/14/17, A/Brisbane/02/016, B/Maryland/15/16, and B/Phuket/3073/13 virus stocks were provided by Novavax, Inc. (Gaithersburg, MD, USA). Histidine-tagged human ACE2 receptor purchased from Sino Biologics (Beijing, CHN). Monoclonal antibody CR3022 [23] was obtained from Creative BioLabs (Shirley, NY, USA, cat # MRO-1214LC). SARS-CoV-2 US-WA recombinant 6-histidine tagged receptor binding domain (RBD) was provided by Novavax, Inc. (Gaithersburg, MD, USA). Histidine tagged B.1.351 spike RBD was obtained from Sino Biologics (cat # 40592-V08H85, Beijing, CHN)

### 2.2. NVX-CoV2373 spike (S) protein and recombinant hemagglutinin vaccines

The SARS-CoV-2 S vaccine was constructed from the full-length; wild-type SARS-CoV-2 S glycoprotein based upon the GenBank gene sequence MN908947 nucleotides 21563-25384. The native, full-length S protein was modified by mutating the putative furin cleavage site (682-RRAR-685 to 682-QQAQ-685) located within the S1/S2 cleavage domain to confer protease resistance. Two proline amino acid substitutions were inserted at positions K986P and V987P (2P) within the heptad repeat 1 (HR1) domain to stabilize SARS-CoV-2 S in a prefusion conformation [18].

The synthetic transgene was codon optimized and engineered into the baculovirus vector for expression in *Spodoptera frugiperda* (Sf9) insect cells (GenScript, Piscataway, NJ, USA). Spike trimers (designated CoV2373) were detergent extracted from the plasma membrane with Tris buffer containing TERGITOL NP-9 detergent and clarified by centrifugation. TMAE anion exchange and lentil lectin affinity chromatography was used to purify S-trimers. Purified CoV2373 was formulated in 25 mM sodium phosphate (pH 7.2), 300 mM NaCl, and 0.02% (v/v) polysorbate [17].

### 2.3. Cloning and expression of hemagglutinin (HA) nanoparticles

Influenza virus A/Kansas/14/17, A/Brisbane/02/016, B/Maryland/15/16, and B/Phuket/3073/13 HA genes were codon optimized for expression in Sf9 insect cells. Synthetic codon optimized HA genes were cloned into pBac1 baculovirus transfer vectors (Millipore Sigma, Billerica, MA, USA). pBac1 plasmids were transfected into Sf9 with Flash-bacGOLD bacmid containing the *Autographa californica* polydedrosis virus genome (Oxford Expression Technology, Oxford UK). Sf9 cells cultures were infected with recombinant baculovirus expressing the HA genes. Recombinant HA was purified as previously described [12].

### 2.4. Animal ethics statement

Noble Life Sciences performed the ferret (*Musteia putorius furo*) immunogenicity study (Sykesville, MD, USA). BioQual, Inc. (Rockville, MD, USA) performed the hamster (*Mesocricetus auratus*) challenge study in an ABSL3 containment facility. Ferrets and Golden Syrian hamsters were maintained and treated according to Animal Welfare Act Regulations, the US Public Health Service Office of Laboratory Animal Welfare Policy on Humane Care and Use of Laboratory Animals, Guide for Care and Use of Laboratory Animals (Institute of Laboratory Animal Resources, Commission on Life Sciences, National Research Council, 1996), an AAALAC accreditation. The studies were conducted in accordance with each institutes’ IACUC approved protocol.

### 2.5. Study design

#### Ferret immunogenicity

Ferrets (n = 30, 15 males and 15 females) were randomized into 5 groups. Animals were immunized by IM with 15 μg or 60 μg HA/strain with without or mixed with 5 μg CoV2373. A comparator group was immunized with 5 μg CoV2373. All vaccines were adjuvanted with 50 μg Matrix-M. All groups were immunized with a prime/boost regimen spaced 21 days apart. Sera were collected for analysis 21 days after the priming dose and 14 days after the booster immunization.

#### Hamster immunogenicity and SARS-CoV-2 challenge

Hamsters (n = 54, 27 males and 27 females) 6-9 weeks old and weighing approximately 100 grams, were randomized into 10 groups (n = 5-6/group). Animals were immunized by intramuscular (IM) injection with 10 μg HA/strain combined with 5 μg or 1 μg CoV2373; 2.5 μg HA/strain combined with 5 μg or 1 μg CoV2373; 10 μg HA/strain; 2.5 μg HA/strain; 5 μg CoV2373; or 1 μg CoV2373. Vaccines were mixed with 15 μg Matrix-M on the day of injection. All groups were immunized with a prime/boost regimen spaced 14 days apart. A placebo group (n = 5) received formulation buffer on days 0 and 14. A sham control group (n = 5) was not immunized. Fourteen days after the first immunization and 14 days after the booster immunization sera were collected for analysis.

### 2.6. SARS-CoV-2 intranasal challenge

Three weeks after the second immunization (study day 35) vaccinated and placebo animals were sedated with 80 mg/kg ketamine and 5 mg kg-1 xylazine in 20 μL sterile phosphate buffered saline (PBS), and inoculated by the intranasal route with 2.0 × 10^4^ pfu of SARS-CoV-2 (strain 2019-nCoV/USA-WA1/2020).

### 2.7. Anti-spike IgG ELISA

Spike protein ELISA was used to determine anti-SARS-CoV-2 spike (S) protein IgG titers in sera. Microtiter plates (Thermo Fisher Scientific, Rochester, NY, USA) were coated with 1.0 µg mL^-1^ of SARS-CoV-2 S protein (CoV2373, Lot# 20Jun20, Novavax, Inc. Gaithersburg, MD, USA). Plates washed with PBS-Tween (PBS-T) and non-specific binding was blocked with TBS Startblock blocking buffer (Thermo Fisher Scientific). Serum samples were serially diluted 3-fold starting with a 1:50 dilution and added to the coated plates, followed by incubation at room temperature for 2 hours. Following incubation, plates were washed with PBS-T and horseradish peroxidase (HRP)-conjugated goat anti-hamster IgG (Southern Biotech, Birmingham, AL, USA) or goat anti-ferret IgG (Abcam, Cambridge, MA, USA) added for 1 hour. Plates were washed with PBS-T and 3,3’,5,5’-tetramethylbenzidine (TMB) peroxidase substrate (Sigma, St Louis, MO, USA) added. Reactions were stopped with TMB stop solution (ScyTek Laboratories, Inc. Logan, UT). Plates were read at OD 450 nm with a SpectraMax Plus plate reader (Molecular Devices, Sunnyvale, CA, USA). EC50 values were calculated by 4-parameter fitting using SoftMax Pro 6.5.1 GxP software. Individual animal anti-spike IgG titers were determined and the group geometric mean titers (GMTs), and 95% confidence intervals (± 95% CI) plotted using GraphPad Prism 8 software. A titer below the assay lower limit of detection (LOD) of 100 (starting dilution) was reported and a value of “50” assigned to the sample to calculate the group GMT.

### 2.8. Human ACE2 receptor blocking antibody ELISA

hACE2 receptor blocking antibody titers were determined by ELISA. Microtiter plates were coated with 1.0 μg mL^-1^ SARS-CoV-2 S protein (CoV2373, Lot# 20Jun20, Novavax, Inc., Gaithersburg, MD, USA) overnight at 4°C. Serum was serially diluted 2-fold starting with a 1:20 dilution and were added to coated wells and incubated for 1 hour at room temperature. After washing, 30 ng mL^-1^ of histidine-tagged hACE2 (Sino Biologics, Beijing, CN) was added to wells for 1 hour at room temperature. HRP-conjugated anti-histidine IgG was added and incubated for 1 hour followed by addition of TMB substrate. Plates were read at OD 450 nm with a SpectraMax Plus plate reader (Molecular Devices, Sunnyvale, CA, USA) and data analyzed with SoftMax Pro 6.5.1 GxP software. The % inhibition for each dilution for each sample was calculated using the following equation in the SoftMax Pro program: 100-[(MeanResults/ControlValue@PositiveControl)*100].

Serum dilution versus % inhibition plot was generated, and curve fitting was performed by 4-parameter logistic (4PL) curve fitting to data. Serum antibody titer at 50% inhibition (IC50) of hACE2 to SARS-CoV-2 rS protein was determined in the SoftMax Pro program. Individual animal hACE2 receptor inhibiting titers, group GMT ± 95% CI were plotted using GraphPad Prism 8 software. For a titer below the assay lower limit of detection (LOD), a titer of < 20 (starting dilution) was reported and a value of “20” assigned to the sample to calculate the group GMT.

### 2.9. Biolayer interferometry competitive binding assay

Competition binning assay was performed using biolayer interferometry (BLI) assay was performed using an Octet QK 384 instrument (FortéBio). BLI studies were done with his_6_-tagged SARS-CoV-2 rS protein receptor binding domain (RBD) coupled to Ni-NTA biosensor tips. Captured RBD is presented in two consecutive steps: 1) experimental serum samples (1:200), pre-immune (day 0) negative control, and positive control prepared with Placebo pool serum (1:200) spiked with 5 μg mL^-1^ of a spike-specific monoclonal antibodies (mAbs); and 2) competing SARS-CoV-2 mAb (5 μg mL^-1^) was loaded onto the RBD biosensor tips and additional binding or competition measured. Data were analyzed using Octet data analysis HT10.0 software and antibody binding matrix was generated. Data were normalized against placebo day 0 serum and percentage of binding and competition of rS RBD mAb to sample was calculated.. Competing antibody concentration (µg mL^-1^) in serum samples were calculated based on percentage of polyclonal antibody competition and concentration of competing mAb.

### 2.10. Hemagglutinin inhibiting antibodies (HAI)

HAI responses against influenza A/Brisbane/02/2018 (H1N1), A/Kansas/14/17 (H3N2), and B strains (B/Maryland/15/16 and B/Phuket/3073/13) were evaluated in serum samples. A 0.75% suspension of human red blood cells (RBC, Biological Specialty Corporation, Allentown, PA, USA) was prepared in Dulbecco’s phosphate-buffered saline (DPBS). Serum samples were treated with receptor-destroying enzyme (RDE) from *Vibrio cholerae* (Denka Seiken, Stamford, TX, USA) at 37°C overnight to eliminate nonspecific red blood cell (RBC) hemagglutinating activity. RDE was inactivated the next day by incubation at 56°C for 1 hour. RDE-treated sera were serially diluted 2-fold in DPBS (starting at 1:10, 25 µL) in 96-well, U bottom plates and incubated with standardized influenza virus concentration (4 HA Units in 25 μL) for 25 minutes. At the end of the incubation, 0.75% suspension of human RBC (50 µL) were added to each well and the plates were incubated at room temperature for 45 minutes.

Hemagglutination inhibition (HAI) was determined by observing the O-ring shape formed by the RBCs in the sample wells and in the negative control wells. The HAI titers were recorded as the reciprocal of the highest serum dilution where HAI was observed (last well with O-ring). For a titer below the assay limit of detection (LOD), a titer of < 10 (starting dilution) was reported and a value “5” assigned to the sample to calculate the group geometric mean titers (GMT).

### 2.11. Micro neutralization (MN) assay

Virus neutralizing antibodies against influenza A/Kansas/14/17 (H3N2), A/Brisbane/02/208 H1N, B/Phuket/3073/13, and B/Maryland/15/2016 were evaluated in serum samples. Serum samples were heat-inactivated at 56 °C for 30Dminutes, 2-fold serially diluted (starting at 1:20, 50 µL) and incubated with 100 TCID_50_ virus (50 µL) for 2 hours. At the end of incubation, 100 µL of 1.5×10^5^ mL^-1^ MDCK cells were added to each well and the plates were incubated with 5% CO_2_ at 37°C for influenza A virus or 32°C for influenza B virus. After 18–22 hours incubation, cells were fixed with 80% cold acetone and incubated with murine monoclonal anti-influenza A or B nucleoprotein (Millipore Billerica, MA, USA) followed by peroxidase-conjugated goat anti-mouse IgG (Kirkegaard and Perry Laboratories, Gaithersburg, MD, USA). Optical density following development with 3-amino-9-ethylcarbazole (AEC) substrate (Sigma Aldrich, Saint Louis, MO, USA) was used to calculate the 50% micro neutralization titer (50% MN) for each serum sample. For a titer below the assay limit of detection (LOD), a titer of < 20 (starting dilution) was reported and a value “10” assigned to the sample to calculate the group geometric mean titers (GMT).

### 2.12. Oral swabs, branchoalveolar lavage (BAL), and lung tissue sample collection

Oral swabs were collected on study days 37, 39, and 42 (2, 4 and 7 dpi). BAL samples and lungs were collected at necropsies carried out on 7 dpi. Lungs were weighed, divided in half; one set was weighed (∼0.1 to 0.2 grams) and snap frozen for virus titer determination. The second set was preserved in formalin for histopathology analysis. For viral load assays, tissues were weighed, placed into pre-labeled Sarstedt cryovials (2/sample), and snap-frozen on dry ice. Lung homogenates were prepared in 0.5 mL RNA-Stat for approximately 20 seconds using a hand-held tissue homogenizer (Omni International, Kennesaw, GA, USA). The samples were clarified by centrifugation and supernatants isolated for viral load determination.

### 2.13. Quantification of subgenomic (sg) RNA by qRT-PCR

The qRT-PCR assay utilized primers and a probe designed to amplify and bind to a conserved region of nucleocapsid (N) gene of coronavirus. The signal was compared to a standard curve and calculated to give copies per mL. For the qRT-PCR assay, viral RNA was extracted from lung homogenates with RNA-STAT 60 (Tel-test”B”) mixed with chloroform, precipitated and suspended in RNase-free 0.04% NaN_3_ (AVE) buffer (Qiagen catalog #1020953). To generate a control for the amplification reaction, RNA was isolated from the virus stock using the same procedure. The amount of viral RNA was determined by comparing it to a known quantity of plasmid control. A final dilution of 10^8^ copies per 3 µL was divided into single use aliquots of 10 µL stored at −80°C. The master mix was prepared with 2.5 mL of 2X buffer containing Taq-polymerase was prepared from the TaqMan RT-PCR kit (Bioline #BIO-78005). From the kit, 50 µL of the RT and 100 µL of RNAse inhibitor added. The primer pair at 2 µM concentration was added in a volume of 1.5 mL. 0.5 mL of water and 350 µL of the probe at a concentration of 2 µM are added and the tube vortexed. For the reactions, 45 µL of the master mix and 5 µL of the sample RNA are added to the wells of a 96-well plate in triplicate.

For control curve preparation, control viral RNA is prepared to contain 10^6^ to 10^7^ copies per 3 µL. Ten-fold serial dilutions of control RNA was prepared using RNAse-free water by adding 5 µL of the control to 45 µL of water and repeating this for 7 dilutions. The standard curve range of 1 to 10^7^ copies/reaction. The sg-N used a known plasmid for its curve. Duplicate samples of each dilution are prepared as described above. For amplification, the plate was placed in an Applied Biosystems 7500 Sequence detector (ThermoFisher Scientific) and amplified using the following program: 48°C for 30 minutes, 95°C for 10 minutes followed by 40 cycles of 95°C for 15 seconds, and 1 minute at 55°C. The number of copies of RNA per mL was calculated by extrapolation from the standard curve and multiplying by the reciprocal of 0.2 ml extraction volume. A range of 50 to 5 × 10^8^ RNA copies per mL for nasal washes. Lung virus load was reported as RNA copies per gram homogenate.

2019-nCoV_N1-F: 5’-GAC CCC AAA ATC AGC GAA AT-3’

2019-nCoV_N1-R: 5’-TCT GGT TAC TGC CAG TTG AAT CTG-3’

2019-nCoV_N1-P: 5’-6-FAM/ACCCCGCAT/ZEN/TACGTTTGGTGGACC/3IABkFQ

Sg-N-F 5’-CGATCTCTTGTAGATCTGTTCTC-3’

Sg-N-R 5’-GGTGAACCAAGACGCAGTAT-3’

Sg-N-P 5’-6-FAM/TAACCAGAA/ZEN/TGGAGAACGCAGTGGG/3IABkFQ

### 2.14. Histopathology

Lung samples collected at necropsy 7 days post-challenge were placed in 10% neutral formalin. Fixed tissues were processed and stained with hematoxylin and eosin (H&E) for histological examination. A board-certified pathologist (Experimental Pathology Laboratories, Inc. (EPI, Sterling, VA, USA) examined H&E slides in a blinded fashion.

### 2.15. Statistical analysis

GraphPad Prism 7.05 software was used for statistical analysis. Serum antibody titers were graphically displayed for individual animals and the geometric mean titer (GMT) and 95% confidence interval (95% CI) intervals plotted. Virus loads were plotted as the median value, interquartile range, and the minimum and maximum values. Student’s t-test was used to determine differences between paired groups as indicated the figure legends. p ≤ 0.05 was considered significant.

## 3. Results

### 3.1. Immunogenicity qNIV/CoV2373 combination vaccine in ferrets

The immunogenicity of the qNIV/CoV2373 combination was evaluated in ferrets. Groups of male and female animals were immunized with a standard or high dose (15 μg HA/strain or high 60 μg HA/strain, respectively) quadrivalent HA (qNIV) combined with 5 μg CoV2373. Comparator groups were immunized with 15 μg HA/strain, 50 μg HA/strain or with 5 μg monovalent CoV2373 vaccines. All vaccines contained 50 μg Matrix-M adjuvant (**Figure 1A**). Human ACE2 receptor inhibiting antibodies levels produced by the combination vaccines were compared to animals immunized with qNIV and CoV2373 components. Animals immunized with qNIV/CoV2373 had slightly elevated levels of hACE2 receptor blocking antibodies 2 weeks after a single dose (GMT = 34-39), which increased 3.2-7.3-fold (GMT 107-202) 2 weeks following the booster immunization. Human ACE2 inhibiting titers were comparable to animals immunized with 5 μg monovalent CoV2373 (GMT = 290). Animals immunized with qNIV alone had no measurable hACE inhibiting antibodies (**Figure 1B**). Hemagglutination inhibiting (HAI) antibody titers were compared between groups. HAI titers to influenza A and B strains were elevated 2 weeks after a single dose and boosted 2-7-fold at 2 weeks following the booster immunization.

**Figure 1.**
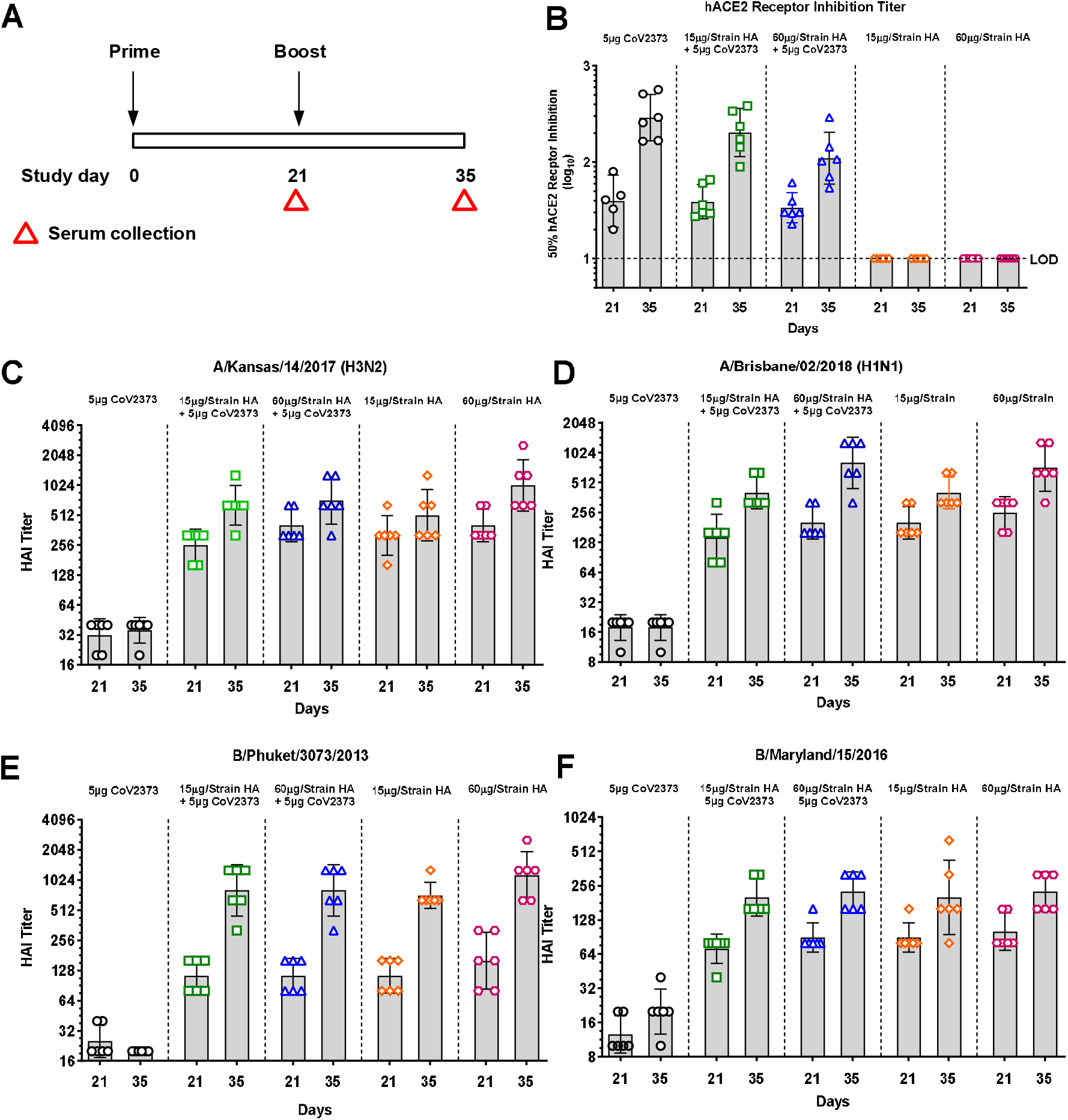
Immune response to SARS-CoV-2 spike glycoprotein and influenza hemagglutinin in ferrets immunized with qNIV/CoV2373 combination vaccine. (**A**) Groups of male and female ferrets (n = 6/group) were immunized with a combination of 15 μg or 60 μg hemagglutinin (HA) per strain with 5 μg CoV2373 with 50 μg Matrix-M. Comparator groups were immunized with 15 μg or 60 μg HA per strain or with 5 μg CoV22373 with 50 μg Matrix-M. All groups were immunized with two doses spaced 21 days apart. Red triangles indicate serum collection days. (**B**) Human angiotensin converting enzyme-2 (hACE2) receptor blocking antibody titer. Hemagglutinin inhibiting antibodies (HAI) titers 21 days after 1-dose and 14 days after the booster immunization (study day 35). (**C**) A/Kansas/14/2017. (**D**) A/Brisbane/02/2018. (**E**) B/Phuket/3070/2013. (**F**) B/Maryland/15/2016. Bars indicate the geometric mean titer (GMT) and error bars indicate the 95% confidence interval.

HAI titers produced by the qNIV/CoV2373 combination were comparable to HAI titers produced by immunization with low or high dose qNIV for all influenza A and B strains. Animals immunized with 5 μg CoV2373 vaccine had no measurable HAI antibodies to A or B strains (**Figure 1C-1F**).

### 3.2. Immunogenicity qNIV/CoV2373 combination vaccine in hamsters

We next evaluated the immunogenicity and protection produced by qNIV/CoV2373 combination vaccine compared to the component vaccines in hamsters challenged with SARS-CoV-2. Groups of hamsters were immunized with qNIV/CoV2373 consisting of 10 μg or 2.5 μg of HA/strain combined with 5 μg or 1 μg CoV2373. Comparator groups were immunized with the qNIV (10 μg or 2.5 μg HA/strain) or with CoV2373 (5 μg or 1 μg). All vaccine formulations were adjuvanted with 15 μg Matrix-M. The placebo group received formulation buffer (**Figure 2A**). Animals receiving the qNIV/CoV2373 combination had elevated anti-S IgG (GMT = 9467 – 25,295) 2 weeks after the first immunization which increased 15-30-fold (GMT = 275,341 – 418,124) 2 weeks following the booster immunization (**Figure 2B, 2C**). Anti-S IgG titers at 2 weeks after the first GMT = 4576 – 43,632) and second doses (GMT = 302,967 – 523,143) of monovalent CoV2373 were comparable to those in animals receiving the combined vaccines (**Figure 2B, 2C**).

**Figure 2.**
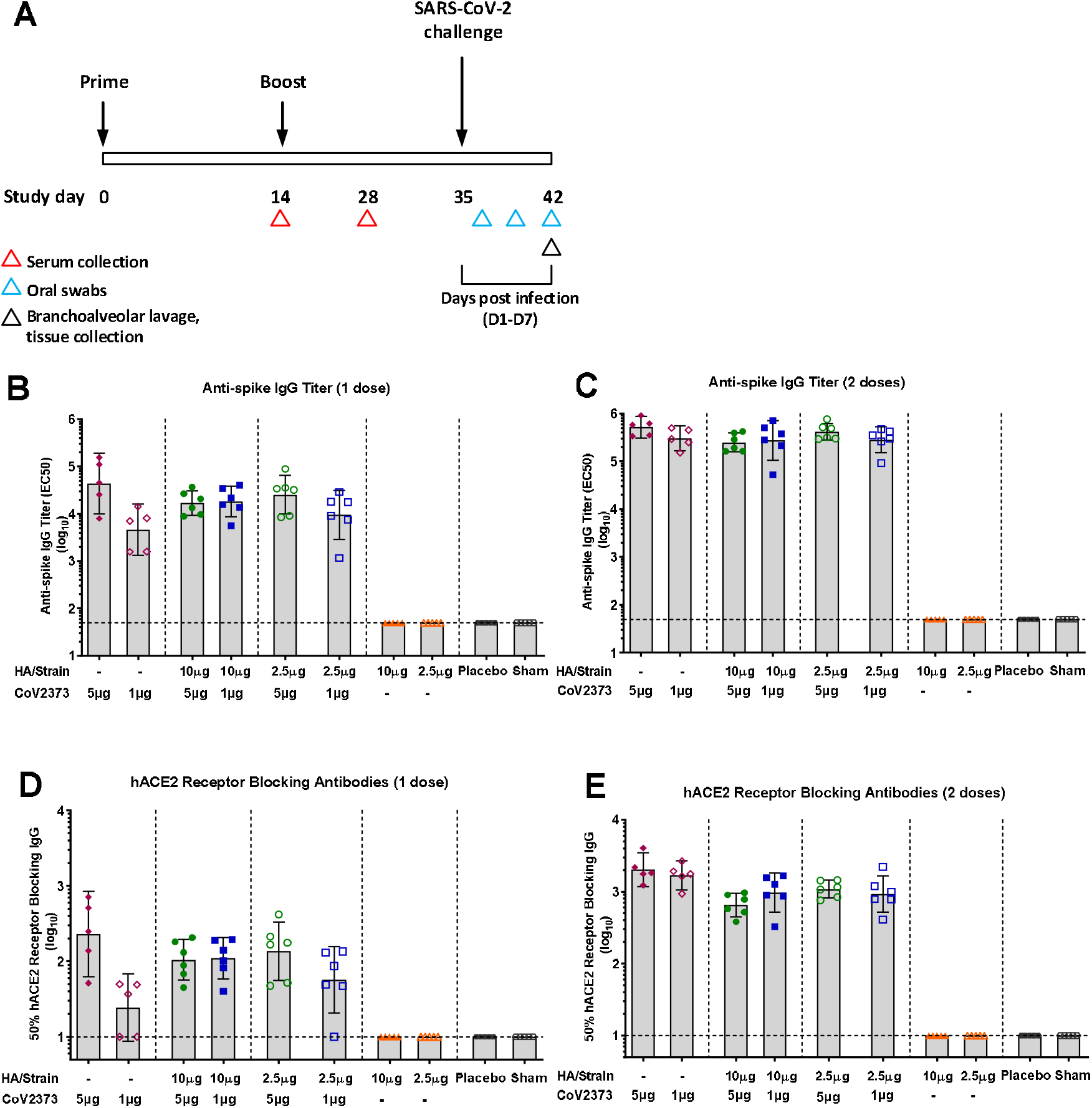
Immune response to SARS-CoV-2 spike glycoprotein and receptor blocking antibodies in hamsters immunized with qNIV/CoV2373 combination vaccine. (**A**) Groups of male and female hamsters (n = 5-6 per group) were immunized with a combination of 2.5 or 10 μg hemagglutinin (HA)/strain with 1 or 5 μg CoV2373 with 15 μg Matrix-M. Comparator groups were immunized with 2.5 or 10 μg HA/strain or 1 or 5 μg CoV2373 with 15 μg Matrix-M. Animals were immunized by the intramuscular route (IM) with 2 doses spaced 14 days apart. The placebo group received formulation buffer. Sera were collected for analysis as indicated by the red triangles. Immunized and placebo animals were infected by the intranasal (IN) route with 2.0 × 10^4^ pfu of SARS-CoV-2 at 21 days after the second immunization. Oral swabs were collected at 2, 4 and 7 days post infection (dpi, blue triangles). Branchoalveolar lavage and lung samples were collected 7 dpi (black triangle). (**B, C**) Anti-spike IgG titer 14 days after 1 dose and 14 days after the booster immunization (study day 28), respectively. (**D, E**) Human ACE2 (hACE2) receptor blocking antibody titer 14 days after 1 dose and 14 days after the booster immunization, respectively. The bars indicate the group geometric mean titer (GMT) and the error bars indicate the 95% confidence interval (95% CI). Individual animal values are indicated by the colored symbols. The horizontal dashed black line indicates the limit of detection (LOD).

Human ACE2 receptor inhibiting antibody levels elicited by qNIV/CoV2373 combination compared to antibody levels elicited by monovalent CoV2373. Hamsters immunized with qNIV/CoV2373 had elevated antibody levels (IC_50_) that block spike binding to the hACE2 receptor after a single dose (GMT = 57 – 136). Receptor inhibiting titers increased 6.2-16.3-fold (GMT = 654 – 1086) following the booster immunization. Human ACE2 inhibiting levels were similar to hamsters following a single immunization with CoV2373 (GMT = 24 – 230). Human ACE2 receptor inhibiting titers increased 7.7-68-fold (GMT = 1636 – 1769) following the booster immunization (**Figure 2D, 2E**).

The immune responses to influenza A and B strains elicited by qNIV/CoV2373 compared to immunization with qNIV. Hamsters immunized with the combination vaccine had high HAI titers to A/Kansas H3N2 (GMT = 113 – 202) and A/Brisbane H1N1 (GMT = 143 - 226) after a single immunization. A/Kansas H3N2 HAI titers increased 6.3 – 14.2-fold (GMT = 1280 – 1810) following the booster immunization and A/Brisbane H1N1 HAI titers increased 5.7 – 10-fold (GMT = 1280-1437) following the booster. (**Figure 3A – 3D**). Animals immunized with qNIV had equivalent HAI titers to the A strains. A/Kansas (GMT = 80-106) and A/Brisbane (GMT = 121-211) after a single immunization. A/Kansas HAI titer increased 24-fold (GMT = 1940-2560) and A/Brisbane HAI titer increased 8–12.1-fold (GMT 1470-1689) after the booster immunization. Animals immunized with the combination vaccine had elevated HAI titers to B/Phuket after as single immunization (GMT = 80-143) that increased 5-9-fold (GMT = 570 – 718) following the second immunization. Animals receiving a single immunization with qNIV had comparable B/Phuket HAI titers (GMT = 735 – 844) after the prime/boost immunization (**Figure 3E, 3F**). Likewise, animals immunized twice with the qNIV/CoV2373 had similar HAI titers to B/Maryland (GMT = 160 - 142), which was equal to HAI titers elicited by immunization with the qNIV (GMT = 106 – 242) (**Figure 3G, 3H**).

**Figure 3.**
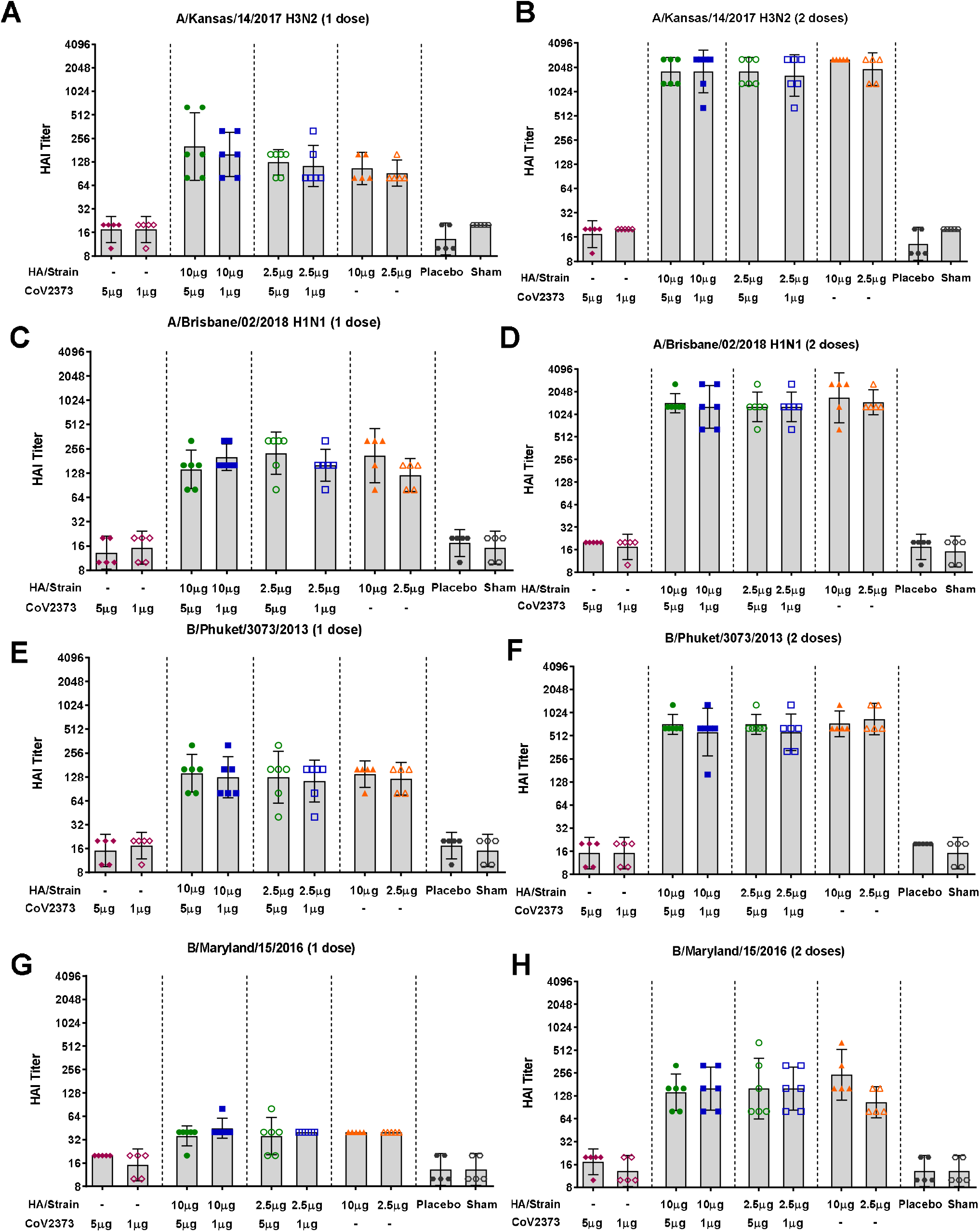
Immune response to influenza HA in hamsters immunized with qNIV/CoV2373 combination vaccine. Groups of hamsters were immunized with the combination of qNIV/CoV2373 vaccine or with the component vaccines as indicated in Figure 2A. Serum was analyzed for hemagglutinin inhibiting antibodies (HAI) titers 14 days after one dose and 14 days after the booster immunization (study day 28). (**A, B**) A/Kansas/14/2017. (**C, D**) A/Brisbane/02/2018. (**E, F**) B/Phuket/3070/2013. (**G, H**) B/Maryland/15/2016. Bars indicate the geometric mean titer and error bars indicate the 95% confidence interval.

Virus neutralizing antibody titers were comparable between groups of hamsters immunized with qNIV/CoV2373 compared to animals immunized with the qNIV alone. Animals immunized with a prime/boost with qNIV/CoV2373 had high neutralizing titers to A/Kansas (GMT = 15,693 – 16,916) and A/Brisbane (GMT = 6992 – 10,507). Immunization with qNIV elicited similar neutralizing titers to A/Kansas (GMT = 13,625 – 19,314) and A/Brisbane (GMT = 20,146 −22,146) (**Figure 4A – 4D**).

**Figure 4.**
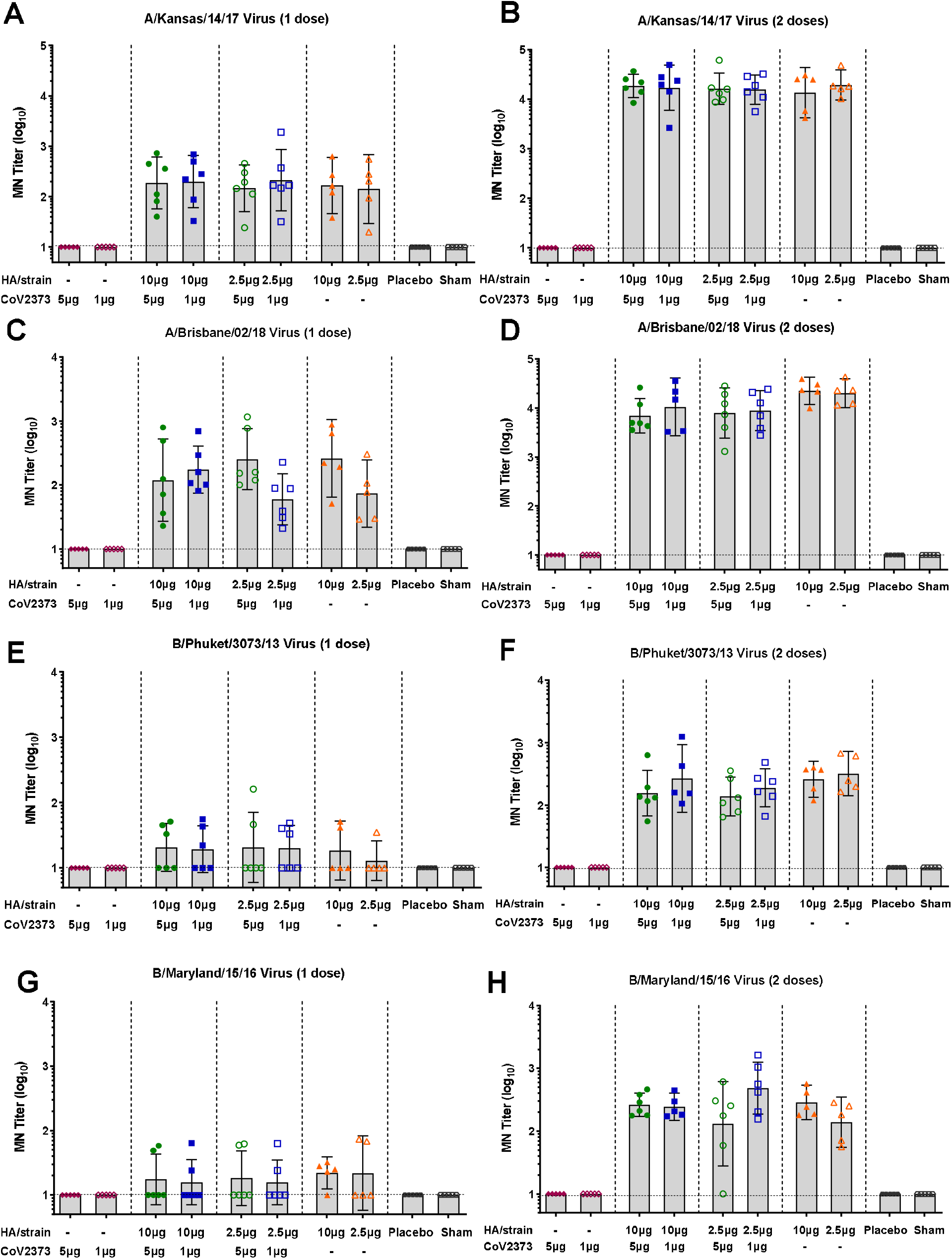
Influenza virus neutralizing antibodies in hamsters immunized with qNIV/CoV2373 combination vaccine. Groups of hamsters were immunized with the combination qNIV/CoV2373 or with the component vaccines as indicated in Figure 2A. Virus neutralizing titers were determined 14 days after 1 dose and 14 days after the booster immunization (study day 28). (**A, B**) A/Kansas/14/2017. (**C, D**) A/Brisbane./02/2018. (**E, F**) B/Phuket/3073/2013. (**G, H**) B/Maryland/15/2016. Bars indicate the geometric mean titer (GMT) and error bars indicate the 95% confidence interval. Horizontal dashed line indicates the limit of detection (LOD).

Similarly, animals immunized with qNIV/CoV2373 or with qNIV had comparable levels of B strain neutralizing titers (**Figure 4E-4H**). Taken as a whole, these results indicate that the qNIV/CoV2373 combination vaccine was immunogenically equivalent to the qNIV and monovalent CoV2373 comparators, indicating there was no interference in the immune responses when qNIV and CoV2373 were co-administered with Matrix-M.

### 3.3. Competitive polyclonal RBD antibodies against neutralizing mAbs

Polyclonal antibodies were induced in hamster competitive with prototype US-WA spike RBD neutralizing mAbs CR3022, NVX.322.3 and NVX.239.12 (**Supplementary Table S1**) regardless of vaccination with monovalent CoV2373 or combination qNIV. Hamster immunized with monovalent CoV2373 had significantly higher polyclonal antibodies competitive against the US-WA RBD mAbs when immunized with 5μg versus 1μg monovalent CoV2373 (**Figure 5 A-C**). The combination of qNIV with 1 μg or 5 μg of CoV2373 rS resulted in some reduction in the levels of competitive polyclonal antibodies measured against the US-WA RBD. A similar pattern of competitive polyclonal antibody responses against the B.1.351 variant were induced in hamsters immunized with monovalent CoV2373 and qNIV/CoV2373 combination vaccines. However, monovalent CoV2373 or the combination vaccine with qNIV induced significant polyclonal antibodies competitive with CR3002 and NVX.322.3 (**Figure 5 D-E**). NVX-239.12 did not bind by to the B.1.351 variant RBD.

**Figure 5.**
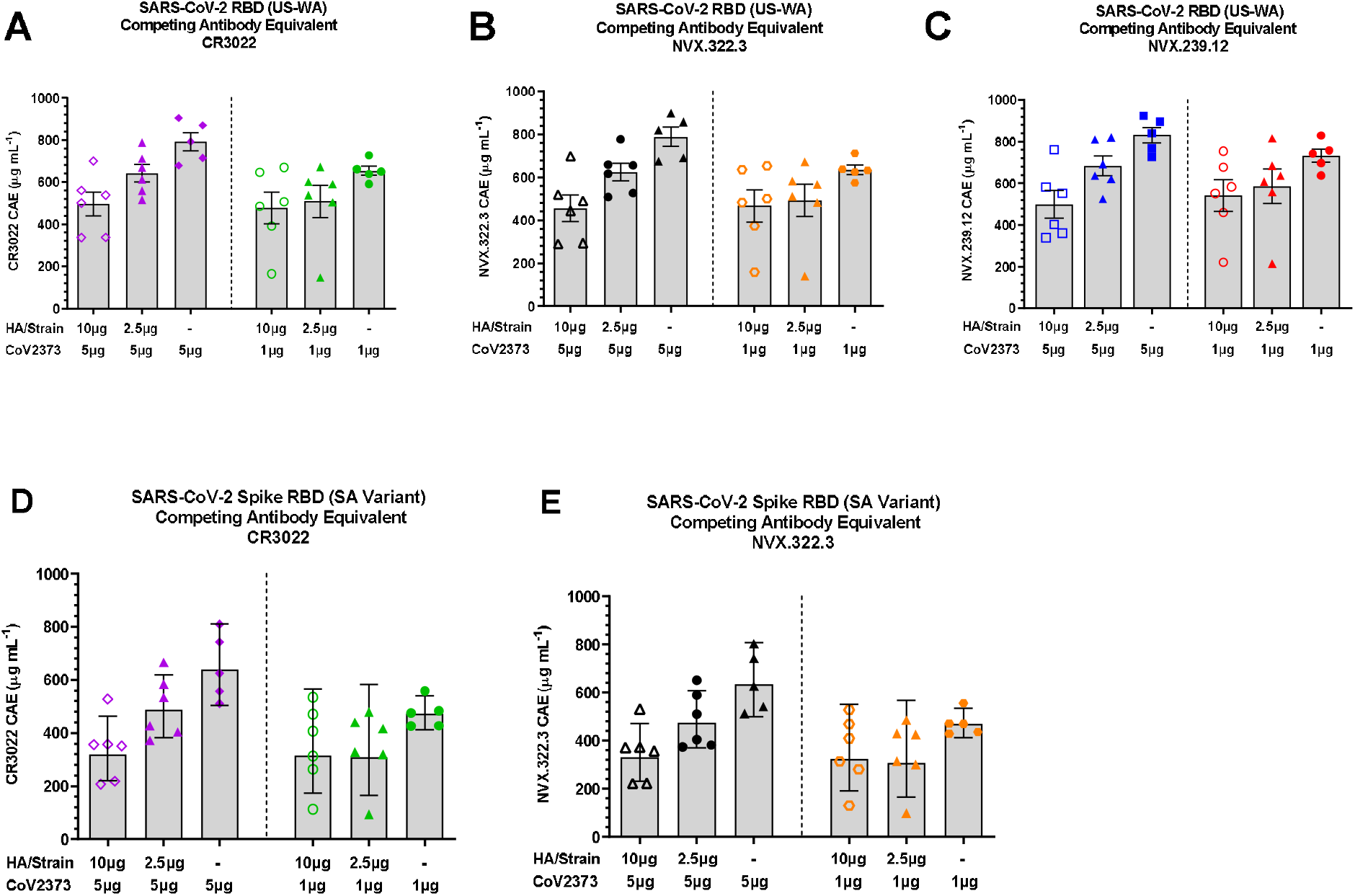
qNIV/CoV2373 vaccine elicits antibody that bind highly conserved cryptic epitopes in the receptor-binding domain (RBD) of SARS-CoV-2 determined by biolayer interferometry (BLI). The specificity of antibodies elicited by qNIV/CoV2373 combination or monovalent CoV2373 was determined by competitive antibody binning of immune serum with receptor site-specific neutralizing monoclonal antibodies by BLI. Horizontal bars indicate the group geometric mean and the error bars indicate the 95% confidence interval. Colored symbols indicate Individual animal values. (**A, B, C**) SARS-CoV-2 US-WA RBD. (**D, E**) SARS-CoV-2 B.1.351 South Africa RBD.

### 3.4. Post challenge clinical signals

To assess the protective efficacy of the combination vaccine, immunized and placebo treated hamsters were challenged with SARS-CoV-2 by the intranasal route 21 days after the second immunization (study day 35) by the intranasal route with SARS-CoV-2. All animals survived the post challenge phase until the scheduled necropsy (7 dpi). Animal weights were monitored daily throughout the post challenge period. Animals receiving the placebo or immunized with the qNIV (2.5 μg or 10 μg HA/strain) lost 12.5% to 15% body weight by 7 dpi. In contrast, animals immunized with 1 μg or 5μg CoV2373 retained their weight with a 2.6% to 5% gain in weight at 7 dpi (**Figure 6A**). Animals immunized with qNIV/CoV2373 retained their weight with a 2.5% to 5% gain in weight at 7 dpi, which was the same as the non-infected sham group (**Figure 6B**).

**Figure 6.**
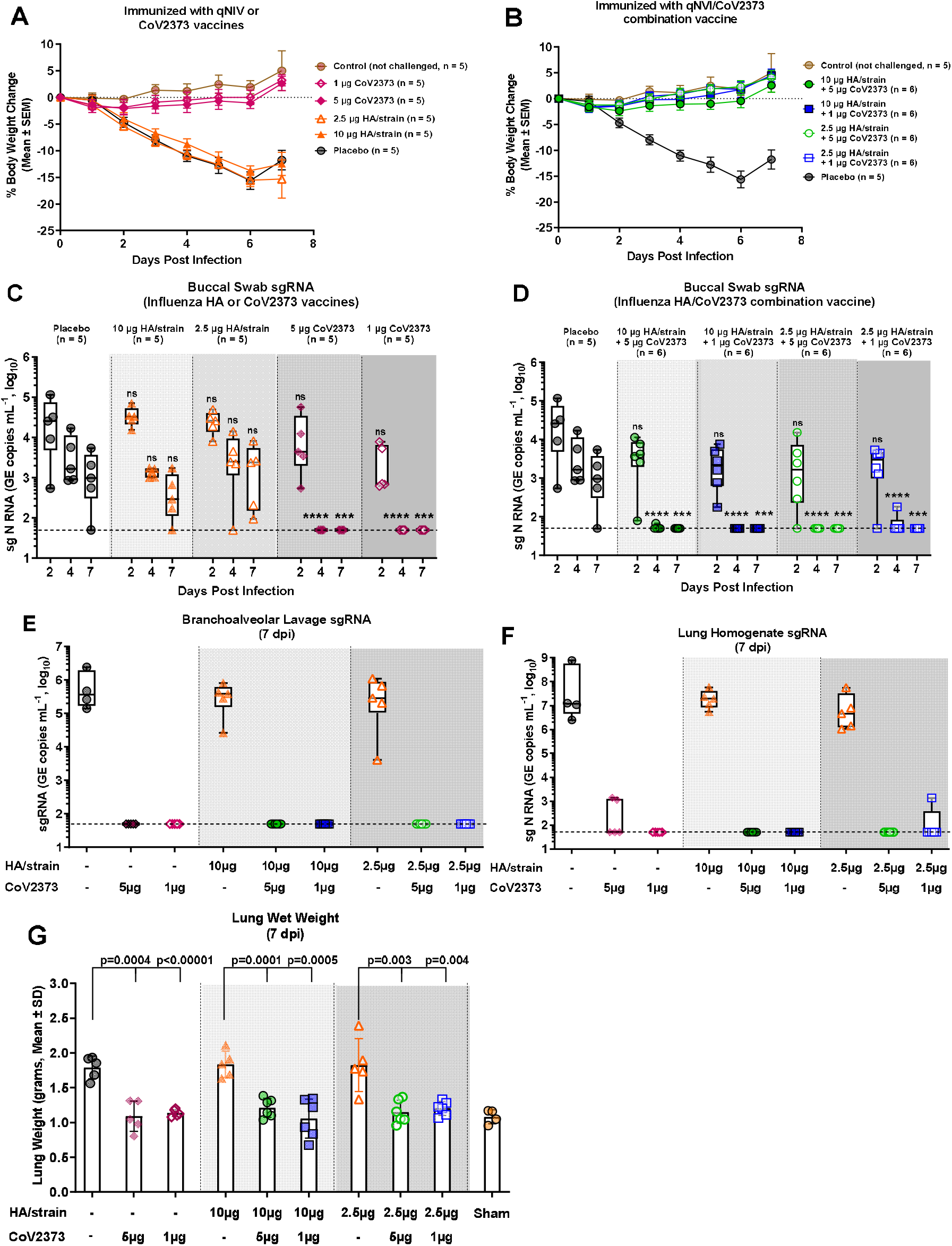
Weight change and protection against upper and lower respiratory infection with SARS-CoV-2 in hamsters immunized with qNIV/CoV2373 combination vaccine. Groups of male and female hamsters (n = 5-6/group) were immunized with a prime/boost regimen of qNIV/CoV2373 or the component vaccines spaced 14 days apart. Three weeks after the second immunization (study day 35), animals were challenge by the intranasal route with 2.0 × 10^4^ pfu. (**A, B**) Change in body weight (percent change following SARS-CoV-2 challenge) was monitored daily for up to 7 dpi. Data are the mean ± SEM for the vaccinated, placebo, and sham groups. (**C, D**) Buccal swabs were taken at 2, 4, and 7 days post infection (dpi) and analyzed for subgenomic (sg) RNA by qRT-PCR. (**E**) sgRNA virus load in branchoalveolar lavage fluids collected 7 days post infection (dpi). (**F**) sgRNA virus load in lung homogenates collected 7 dpi. In the box-and-whisker plots, the median is indicated by the horizontal line, the top and bottom of the box indicates the interquartile range, and the whiskers indicate minima and maxima for each experimental group (n = 5-6/group). Individual animal values are indicated by colored symbols. The dashed line indicates the assay limit of detection (LOD). Student’s t-test (paired, two tail) was used to determine significant differences in levels of viral sgRNA between placebo treated animals and immunized animals a 2, 4 and 7 dpi. Not significant (ns), ***p≤ 0.001, ****p ≤ 0.0001. (**G**) Weight of lungs collected at 7 dpi from vaccinated, placebo treated, and untreated sham animals. The bars indicate the mean and the error bars indicate the ± standard deviation (SD). Individual animal values are indicated by the colored symbols. Student’s t-test (paired, two tail) was used to determine significant differences in lung weigh between the indicated paired groups.

### 3.5. Subgenomic virus mRNA in oral swabs, bronchoalveolar lavage (BAL), and lung samples

To examine the efficacy of the qNIV/CoV2373 vaccine, virus load in the upper and lower respiratory tract was determine using qRT-PCR designed to detect SARS-CoV-2 subgenomic (sg) SARS-CoV-2 nucleocapsid (N) RNA. Oral swabs were collected 2, 4 and 7 days post infection (dpi). The highest levels of sgRNA were observed in oral swabs of placebo treated animals with a median peak of 4.4 (range 2.7-5.1) log_10_ RNA copies mL^-1^ at 2 dpi. Viral levels remained elevated at 3.2 (range 2.9-4.2) log_10_ RNA copies mL^-1^ at 4 dpi and declined to 3.0 log_10_ (range 1.7-3.7) RNA copies mL^-1^ at 7 dpi. Viral RNA levels were not significantly different in oral swabs from animals immunized with 10 μg or 2.5 μg HA/strain with the highest levels of sgRNA of 4.4-4.5 (range 3.9-4.9) log_10_ RNA copies mL^-1^ at 2 dpi, 3.2-3.4 (range 1.7-3.7) log_10_ RNA copies mL^-1^ at 4 dpi, and 2.5-3.4 (1.7-3.9) log_10_ RNA copies mL^-1^ at 7 dpi. Oral swabs from animals immunized with 5 μg or 1 μg CoV2373 had detectable viral RNA at 2 dpi (2.9-3.6 log_10_ copies mL^-1^). No viral RNA was detected in swabs of animals immunized with CoV2373 at 4 or 7 dpi (**Figure 6C**). Viral RNA (3.2-3.5 log_10_ RNA copies mL^-1^) was detected only in swabs collected 2 dpi in animals immunized with qNIV/CoV2373, while no replicating virus was detected at 4 or 7 dpi (**Figure 6D**).

BAL washes were obtained at necropsy and analyzed for viral RNA. The placebo group and the groups immunized with 10 μg or 2.5 μg HA/strain had the highest median levels of viral RNA. Aspirates from the placebo group had the levels of replicating virus with a median of 5.6 (range 5.1-6.4) log_10_ sgRNA copies mL^-1^. Virus load in BAL was also high in animals immunized with 10 μg HA/strain with a median of 5.6 (range 4.4-5.9) log_10_ RNA copies mL^-1^ and 2.5 μg HA/strain with a median of 5.5 (range 3.6-6.0) log_10_ RNA copies mL^-1^. Little or no viral RNA was detected in BAL samples obtained from animals immunized with CoV2373 (1 μg or 5 μg) or with the qNIV/CoV2373 combination (**Figure 6E**).

Lung tissues collected at necropsy and were evaluated for virus load. Lung homogenates from placebo treated and animals receiving qNIV had the highest virus load: placebo median 7.1 (range 6.4-8.9) log_10_ RNA copies gram^-1^; 10 μg HA/strain 7.3 (range 6.7-7.8) log_10_ RNA copies gram^-1^; and 2.5 μg HA/strain 6.7 (range 6.0-7.7) log_10_ RNA copies gram^-1^. Lung homogenates from animals immunized with CoV2373 or the qNIV/CoV2373 combination had little or no detectable virus (**Figure 6F**).

### 3.6. Macroscopic and microscopic observations

All animals survived until the scheduled necropsy (study day 42). Lungs were collected and weighed. Lung weights were significantly higher (p ≤ 0.003) in animals treated with placebo or immunized with qNIV compared to animals immunized with CoV2373 or qNIV/CoV2373 combination (**Figure 6G**). Microscopic findings in the airways of placebo treated and qNIV immunized animals were the same and consisted of bronchoalveolar hyperplasia, mixed alveolar inflammation, perivascular inflammation with edema in the surrounding tissues, and rare syncytial cells. There were no remarkable findings in the lungs of animals immunized with CoV2373 or qNIV/CoV2373 vaccines (**Figure 7A-7J**). These results are consistent with high virus load and pneumonia in animals treated with placebo and receiving qNIV, while lungs of CoV2373 and qNIV/CoV2373 vaccinated animals were normal and free of virus.

**Figure 7.**
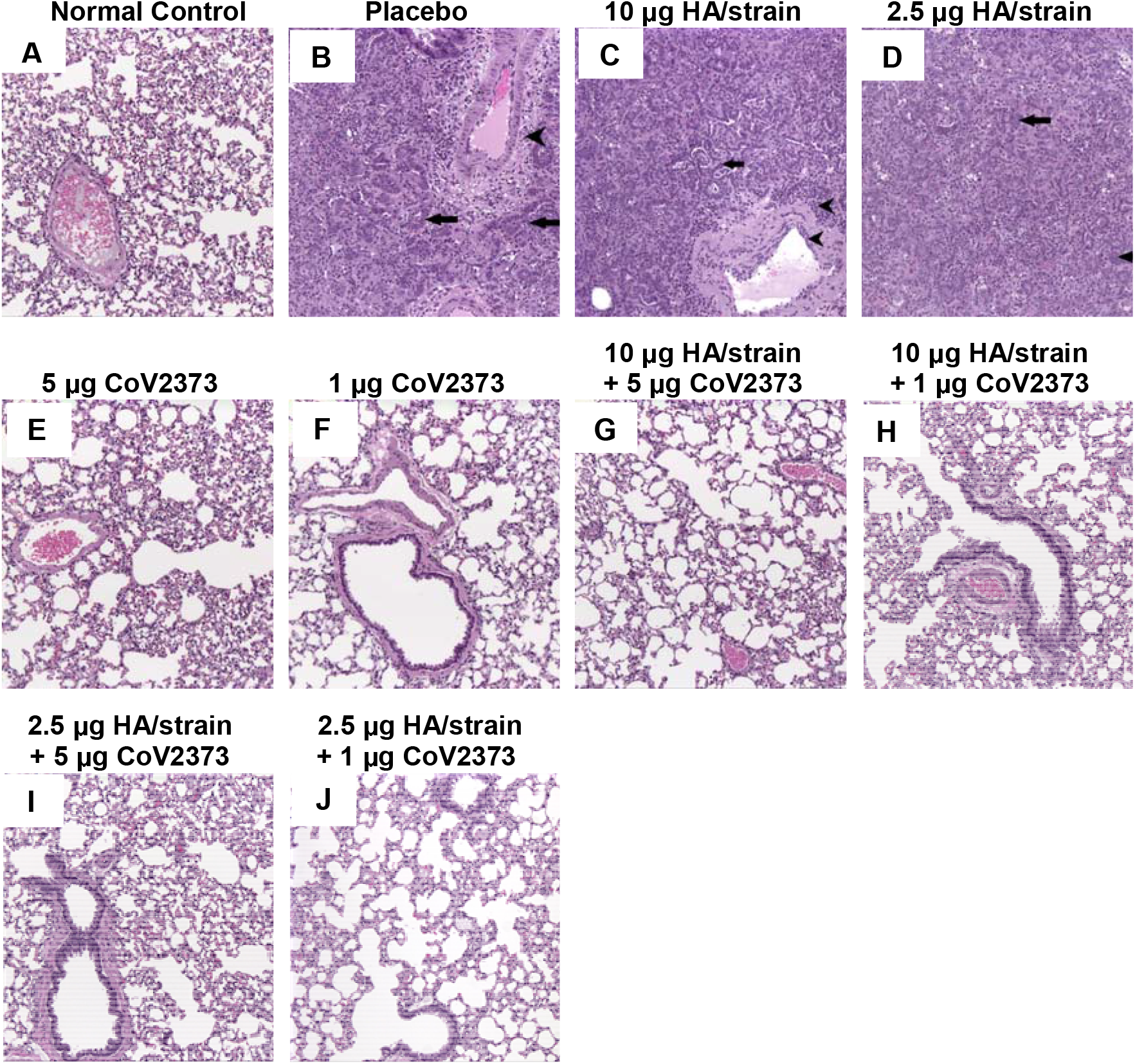
Histopathologic changes in lungs of hamsters immunized with qNIV/CoV2373 combination vaccine and challenged with SARS-CoV-2. Groups of male and female hamsters were immunized with the combination qNIV/CoV2373 with 2.5 μg or 10 μg hemagglutinin (HA) per strain combined with 1 μg or 5 μg CoV2373 adjuvanted with Matrix-M. Comparator groups were immunize with 2.5 μg or 10 μg HA/strain or with 1μg or 5μg CoV2373 recombinant spike adjuvanted with 15 μg Matrix-M. All groups were immunized with 2 doses placed 14 days apart. The placebo group received formulation buffer. Three weeks following the second immunization (study day 35) all animals were challenged intranasally with 2.0 × 10^4^ pfu SARS-CoV-2 and lungs collected 7 dpi. Histologic images were stained with hematoxylin and eosin (H&E). (**A**) Sham control image shows normal lung. (**B**) Microscopic findings in the lungs of placebo treated animals showed the airways were consolidated by bronchiolo-alveolar hyperplasia (arrows) with mixed alveolar inflammation. Mononuclear inflammatory cells surround blood vessels (arrowhead) with edema expanding surrounding tissues (arrowhead). (**C, D**) Microscopic findings in the lungs of animals immunized with qNIV show histologic changes similar to the placebo group including extensive bronchiolo-alveolar hyperplasia (arrows) with mixed alveolar inflammation with perivascular inflammation and expanded vessel walls (arrowheads). (**E, F**) Images of lungs from animals immunized with monovalent CoV2373 show no significant microscopic findings. (**G, H, I, J**) Images of lungs from animals immunized with qNIV/CoV2373 combination vaccine show no significant microscopic findings. Magnification 10X.

## Discussion

We describe herein the development of a novel combination respiratory vaccine containing quadrivalent full-length HA nanoparticles formulated with full-length, prefusion stabilized SARS-CoV-2 spike protein nanoparticles, together adjuvanted with Matrix-M. Both Matrix-M adjuvanted quadrivalent seasonal influenza HA qNIV vaccine alone and the Matrix-M adjuvanted SARS-CoV-2 spike protein vaccine NVX-CoV2373 alone were successfully evaluated for safety and induction of protective responses in pivotal Phase 3 clinical trials [9-11]. The confluence of the COVID 19 pandemic and the recent generation of pivotal data from influenza qNIV vaccine has created a unique opportunity to advance the development of a novel combination-respiratory vaccine.

Previous combination vaccines have almost exclusively been developed for pediatric applications. In the process of development of a combination vaccine, each component or pathogen specific components should have been shown in pivotal trials to modulate disease or induce relevant immunity considered protective. This provides regulators with assurance that when combined and evaluated with similar immune measures considered correlative with protection, each component is likely to be effective. The combination respiratory vaccine consists of a recombinant hemagglutinin quadrivalvent nanoparticle influenza vaccine (qNIV) formulated together with a recombinant SARS-Cov-2 spike protein have met similar criteria. The previous development of the individual components of the combination respiratory vaccine therefore, has derisked the approval pathway, as the HA component induced levels of immunity that are deemed reasonably likely to be protective in a pivotal trial or and the rS vaccine has repeatedly demonstrated efficacy.

An additional needed capability for a successful combination respiratory vaccine will be to provide coverage in the face of the inevitable viral evolution. The RNA respiratory viruses are especially prone to rapid evolution, often under immune pressure from the host and antigenic shift from zoonotic sources. Evolution of the SARS-CoV-2 virus under immune pressure in South Africa has led to apparent outbreaks in populations where some level of herd immunity had been established [10]. In the context of seasonal influenza, viral evolution is a key challenge for effective immunization. Recently, multiple severe A(H3N2)-predominant influenza seasons occurred in the face of recurrent reports of poor field vaccine effectiveness [24]. This appears driven by both antigenic mismatches arising from egg-based vaccines and the viruses themselves because of a rapid rate of antigenic evolution,

We recently described immune responses to conserved, broadly neutralizing epitopes using Matrix-M adjuvanted recombinant hemagglutinin (HA) trivalent nanoparticle influenza vaccine (tNIV) [13, 16]. The full-length surface glycoprotein retains fidelity to the wild-type structure of circulating virus HA sequences but provides for presentation of otherwise immunologically cryptic epitopes that stimulate broadly neutralizing antibodies [13]. In clinical studies we demonstrated improved induction of wild-type hemagglutination inhibition antibody titers against A/H3N2 drift variants isolated over several years when compared with egg-derived, trivalent high-dose inactivated influenza vaccine (IIV3-HD) [15].

In this study, we demonstrate that a combination of qNIV/CoV2373 vaccine induced competitive polyclonal antibody responses against not only US-WA also the B.1.351 South Africa variant of SARS-CoV-2 spike protein RBD neutralizing epitopes. Demonstrating the potential for neutralizing epitopes that are in common between US-WA and B.1.351 RBD and induced by CoV2373 vaccine. Nanoparticle vaccines have been shown to induce responses against otherwise cryptic or hidden epitopes [14]. In the phase 2b clinical trial with CoV2373 vaccine may have also induced antibodies against protective epitopes not seen in natural infection [10].

The remarkable events of 2019-20 saw both the appearance and surge of a novel coronavirus and COVID-19 disease concomitant with a near absence of seasonal influenza. At this time, it has been proposed that COVID-19 control measures may have led to the diminution of seasonal influenza cases while others have proposed ecological mechanisms [25]. In either case, the past centuries indicated that influenza would recirculate, and cause disease and highly infectious and clinically important SARS-CoV-2 and emerging variants are likely to continue to evolve globally. Co-infection of influenza and SAR-CoV-2 has been described [26]. Common clinical symptoms of COVID-19 include fever, chills, cough, and dyspnea, making it difficult to diagnose influenza virus infection. A retrospective study of hospitalized patients positive for SARS-CoV-2 with severe disease showed 12% (64 of 544 patients) were co-infected with influenza A (84%, 54 or 64) and influenza B (16%, 10 of 64) [27]. The clinical impact of influenza and SARS-CoV-2 co-infection, however, is unknown [28].

The future need for seasonal influenza and the ongoing need for SARS-CoV-2 vaccines makes the prospect of a combination vaccine highly desired due to the logistics of immunization with two vaccines annually. In this report, we describe a co-formulated influenza and SARS-Cov-2 nanoparticle vaccine to broadly protect against co-circulating seasonal influenza and COVID-19 viruses and with a potential due to broadly protective responses to address the challenge of emerging influenza strains and SARS-CoV-2 variants.

## Acknowledgements

The authors wish to recognize the contributions of Dr. Swagata Kar, Bioqual Study Director for conduct of the study and Dr. Shannon Wallace, Experimental Pathology Laboratories evaluated lung tissues for histopathology changes associated with virus challenge.

## Author contributions

NP, BZ, MGX, JHT, ADP, MJM, VS, LF, and GS contributed to conceptualization of experiments, generation of data and analysis, and interpretation of the results. NP, JHT, SM, RF and BZ performed experiments. MGX, NP, and MJM coordinated projects. GS, GG, MGX, NP, VS, LF and LE contributed to drafting and making critical revisions with the help of others.

## S1. Supplementary Information

## S1. Methods and Description

### S1.1 SARS-CoV-2 spike monoclonal antibodies

SARS-CoV-2 spike protein-specific mouse monoclonal antibodies were generated using modified standard method [1]. Briefly, female BALB/c mice were immunized with SARS-CoV-2 US-WA recombinant spike (rS) protein adjuvanted with Matrix M1 on day 0 and day 14 by intramuscular injection. A fusion boost by intraperitoneal injection without adjuvant was given 4 days before spleens were collected and homogenized into single cell suspensionsA magnetic cell sorting system (Miltenyi Biotec, Auburn, CA, USA) was used to deplete splenocyte suspension of IgM B cells. IgG enriched splenocytes were fused P3X63.Ag.6.5.3 myeloma cells [1]. Hybridomas were screened by a SARS-CoV-2 rS ELISA and positive cultures subcloned by limiting dilution. Clones were expanded in serum free media and antibodies purified with a protein-G affinity column. Spike receptor binding domain (RBD) specific monoclonal antibodies NVX.239.12 and NVX.322.3 are described in Table S1.

[1] Apiratmateekul N, Phunpae P, Kasinrerk W. A modified hybridoma technique for production of monoclonal antibodies having desired isotypes. *Cytotechnology*. 2009. 60:53. https://doi:10.1007/s10616-009-9222-z.

**Table S1.** Functional characteristics of SARS-CoV-2 spike monoclonal antibodies.

| Antibody | Isotype | SARS-CoV-2 spike binding<br>(EC<sub>50</sub>, ng mL<sup>-1</sup>) | SARS-CoV-2 spike RBD<sup>a</sup><br>(EC<sub>50</sub>, ng mL<sup>-1</sup>) | ACE2<sup>b</sup> receptor inhibition<br>(IC<sub>50</sub>, ng mL<sup>-1</sup>) | 100% SARS-CoV-2 neutralization<br>(IC<sub>100</sub>, ng mL<sup>-1</sup>) |
| --- | --- | --- | --- | --- | --- |
| NVX.239.12 | IgG1/k | 7 | 32 | 56 | 12 |
| NVX.322.03 | IgG1/k | 28 | 250 | 275 | 185 |

<sup>a</sup> RBD: receptor binding domain <sup>b</sup> ACE2: angiotensin-converting enzyme 2 receptor

